# A new *in vitro* checkerboard-parasite reduction ratio interaction assay for early de-risk of clinical development of antimalarial combinations

**DOI:** 10.1101/2022.04.19.488858

**Authors:** Sebastian G. Wicha, Annabelle Walz, Mohammed H. Cherkaoui-Rbati, Nils Bundgaard, Karsten Kuritz, Christin Gumpp, Nathalie Gobeau, Jörg Möhrle, Matthias Rottmann, Claudia Demarta-Gatsi

## Abstract

The development and spread of drug resistant phenotypes substantially threaten malaria control efforts. Combination therapies have the potential to minimize the risk of resistance development but require intensive preclinical studies to determine optimal combination and dosing regimens. To support the selection of new combinations, we developed a novel *in vitro-in silico* combination approach to help identify the pharmacodynamic interactions of the two antimalarial drugs which can be plugged into a pharmacokinetic/pharmacodynamic model built with human monotherapies parasitological data to predict the parasitological endpoints of the combination. This allows to optimally select drug combinations and doses for the clinical development of antimalarials. With this assay, we successfully predicted the endpoints of two phase 2 clinical trials in patients with the artefenomel - piperaquine and artefenomel - ferroquine drug combinations. Besides, the predictive performance of our novel *in vitro* model was equivalent to the humanized mouse model outcome. Lastly, our more granular *in vitro* combination assay provided additional insights into the pharmacodynamic drug interactions compared to the *in vivo* systems, e.g. a concentration-dependent change in the E_max_ and the EC_50_ values of piperaquine or artefenomel or a directional reduction of the EC_50_ of ferroquine by artefenomel and a directional reduction of E_max_ of ferroquine by artefenomel. Overall, this novel *in vitro-in silico*-based technology will significantly improve and streamline the economic development of new drug combinations in malaria and potentially also in other therapeutic areas.

## INTRODUCTION

Poverty-associated infectious diseases such as malaria still inflict extensive morbidity and mortality in resource-poor countries. In 2020 there was an estimated 241 million malaria cases and 627,000 deaths worldwide *(1)*. Malaria eradication poses significant intertwined challenges at the logistical, political, and scientific level. The World Health Organization (WHO) recommends the development of combination therapies *(2)* to treat malaria, so that the substances in the combination can rescue each other if resistance to one of them evolves and, together, still provide an adequate clinical and parasitological response (ACPR) of 95%. Given the substantial numbers of possible combinations, the determination of optimal combination and dosing regimens to achieve the target efficacy is complex and requires more sophisticated preclinical assays.

Although several new antimalarial combinations have been developed, numerous of these were discontinued in the clinical phase after efficacy studies in field settings *(3, 4)*, thus, stressing the need for more sophisticated preclinical assays to increase the success rate of novel antimalarial combinations in clinical development. This would allow to focus only on drug combinations that meet all the required criteria and thus accelerate the development and commercialization of new antimalarial combinations. Conventionally, antimalarial monotherapies and recently also antimalarial combinations are evaluated *in vivo* in a severe immunodeficient NSG (NOD/SCID/IL2R_γ_^−/−^) mouse model engrafted with human erythrocytes and infected with an adapted *P. falciparum* 3D7 strain (*Pfalc*EryHuMouse) *(5– 7)*. Another, even more resource intensive setting is the human volunteer infection study (VIS), in which healthy volunteers are infected with *P. falciparum* 3D7 and treated before becoming symptomatic *(8)*. Apart from ethical concerns regarding the use of animals, and the considerable amount of time and cost required by these *in vivo* studies, only a small number of doses can be tested, hence preventing the exploration of the full combined response surface of the antimalarial effects. Another important limitation of these studies is the uncertainty of parasite killing following drug treatment since conventional methods cannot reliably distinguish viable from dying or dead parasites *(9, 10)*. Indeed, the parasite clearance rate (viable versus dead parasites), measured immediately after treatment, may differ between *in vivo* models owing different clearance mechanisms of dead parasites from the bloodstream. These inaccuracies make it difficult to estimate the maximum effect of the antimalarials and blur pharmacokinetic (PK)/pharmacodynamic (PD) calculations.

Historically, *in vitro* growth inhibition assays *(11, 12)*, presented graphically as isobolograms, have been used to evaluate PD drug-drug interactions (DDIs) to inform the development of novel antimalarial combination treatments. These assays aim to determine the drug concentration added to an *in vitro* culture of parasites that reduces their density to 50% of the untreated control, called the 50% inhibitory concentrations (IC_50_). The assessment is made at a single time point, usually 72 hours after the drug has been added to the culture of parasites. For a combination, an isobole showing the Fractional Inhibitory Concentration (FIC) (the ratio of IC_50_ of the combination to the IC_50_ of the monotherapy) for drug A versus the FIC for drug B is plotted. The PD DDI is then classified as additive, synergistic or antagonistic if the shape of the isobole is linear, concave, or convex, respectively. However, these assays are static and do not allow to identify the PK/PD parameters such as E_max_ or EC_50_ in combination. In addition, isobologram results are prone to inconsistencies between individual studies. For example, while the interaction between tafenoquine and chloroquine was found to be antagonistic or additive by Gorka et al. *(13)*,whereas Bray et al. Al. found the interaction to be synergistic *(14)*. Besides, it is difficult to relate an antagonistic or synergistic effect from an isobologram to the parasitological endpoints. Hence, no current method fully meets the requirement for testing new antimalarial combinations in a reasonable time frame against their chance of success to achieve the target of 95% adequate clinical and parasitological response at day 28 (ACPR28) in patients. To overcome these problems, and to optimally select drug combinations and leverage preclinical data to inform first-in-human clinical studies about potential pharmacodynamic DDIs, we have developed a novel *in vitro-in silico-*based combination technology: the checkerboard-Parasite Reduction Ratio (PRR) interaction assay combined with a PK/PD model-based approach (Figure 1). In accordance with the 3R’s principle by replacing and/or reducing the use of animals, this assay aims to minimise the knowledge gap in translational research and clinical development. It is based on the dynamic checkerboard analysis developed for testing antibiotics combinations *(15)* with rationally selected drug concentrations *(16)* and on the *Plasmodium* growth inhibition assay in combination with a PRR assay, the gold standard to evaluate the maximum parasite killing and informs about parasite viability at different time-points after drug exposure *(17)*. Together, this allows to investigate of the effect of several different drug concentrations in monotherapy and/or in combination with a partner drug.

**Fig. 1.**
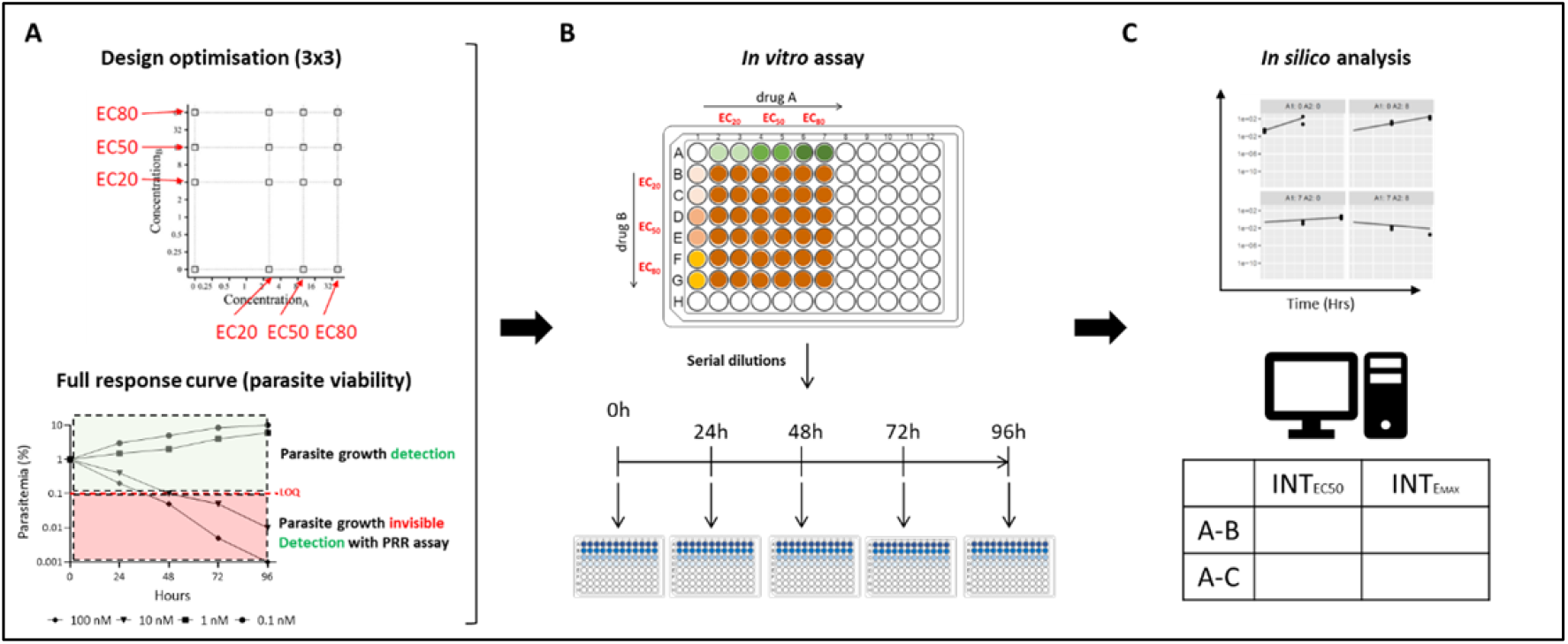
Scheme showing the *in vitro* parasite viability assay for identification of pharmacodynamic antimalarial drug-drug interaction parameters. (A) Schematization of the optimizations made to the assay to provide the relevant data for modeling. The use of effective concentrations (EC20, EC50 and EC80) reduces the number of experiments needed. The parasite reduction ratio (PRR) assay elucidates the full response curve and can also detect differences and interactions on the rate of parasite killing. (B) Parasite killing kinetics were determined by culturing *P. falciparum* parasites in the presence of antimalarial drugs at different concentrations (EC20, EC50, and EC80) as either monotherapy or combinations at different time points. Aliquots corresponding to 1 ml of culture are taken at specific time points, washed and the free parasites cultured with fresh erythrocytes under limiting serial dilution conditions. Parasite growth is subsequently monitored after 21 days and the initial number of viable parasites in the aliquot is calculated (see Material and Methods). (C) Parasite kinetics were modelled with systems of differential equations. Pharmacodynamic drug interactions were quantified using the General Pharmacodynamic Interaction (GPDI) model as shifts of EC50 or Emax. The *in vitro* model parameters were used for clinical trial simulations and compared to real phase 2 clinical data.

The *in vitro* parasite viability data generated with this assay were used in conjunction with state-of-the art PK/PD modelling techniques to describe the killing rate of drugs over time alone and in combination. Here we show how the assay allows to identify the PK/PD relationship as a function of the concentrations of drug A and drug B. We describe the development of this novel approach and its application to two new antimalarial drug combinations, recently evaluated in clinical trials, *i*.*e*. artefenomel (AF) and piperaquine (PQP), as well as AF and ferroquine (FQ). Clinical trial simulations with the *in vitro-in silico* checkerboard-PRR derived PK/PD relationship were performed and predictions were compared with the observations in patients, demonstrating that this hybrid clinical PK-*in vitro* PD interaction model can be exploited for clinical trial simulations of antimalarial drug combinations.

## RESULTS

### *In vitro* checkerboard-PRR interaction assay provides highly granular PD data to successfully quantify complex PD interaction of E_max_ and EC_50_

#### Artefenomel (AF) and piperaquine (PQP) combination

For the *in vitro* checkerboard-PRR interaction assay, concentrations of 0, 7, 12 and 100 nM or 0, 8, 12, 100 nM were selected for AF and PQP, resulting in staggered parasite killing profiles over time. The viable parasite-time course profiles for the AZ – PQP combination were modelled using a time-kill modelling approach based on the *in vitro* checkerboard-PRR interaction assay data (Figure 2). Thereby, the combined drug effects were modelled using the general pharmacodynamic interaction model (GPDI) model *(18)* that provided a flexible framework to quantify PD interactions as shifts of potency (EC_50_) or maximum effect (E_max_) or both at the same time. Moreover, with the GPDI model the directionality of the drug interactions was quantified *i*.*e*. which drug took the role of the perpetrator or victim. The parameter estimates of the PK/PD model are presented in Table S1. Assuming Bliss Independence as underlying additivity criterion, the following PD interactions were estimated quantifying deviations from Bliss Independence:

**Figure 2:**
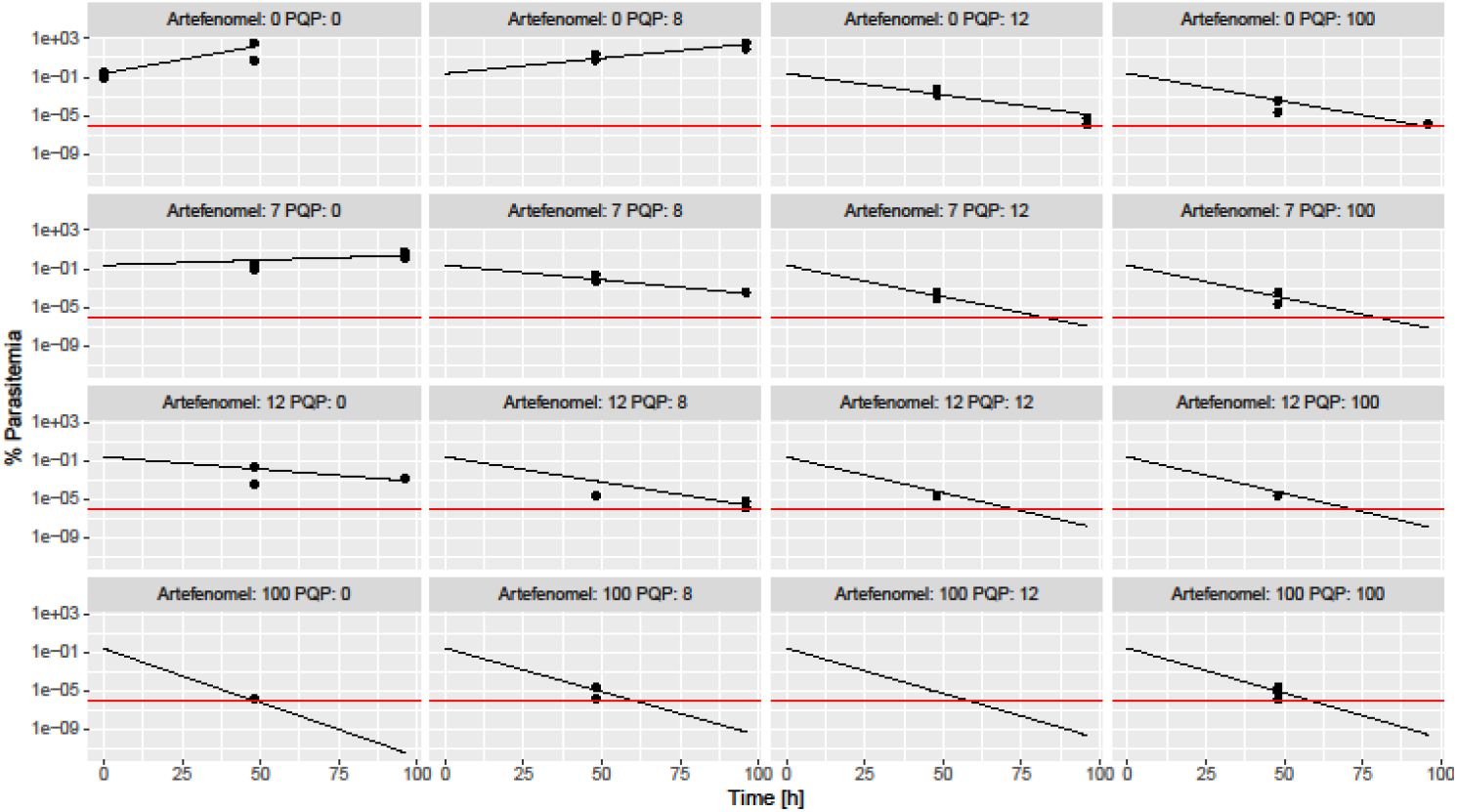
Individual model fits for artefenomel - piperaquine (AF - PQP) for the *in vitro* PRR checkerboard-PRR interaction assay data. Observed parasitaemia (black points) and model-predicted parasitaemia (black lines) time courses at different concentrations of AF and PQP (numbers in facet denote the concentration in nM of each combination partner); red line: quantification limit.

- The EC_50_ of AF was increased 10.7-fold mediated by PQP
- The EC_50_ of PQP was reduced by 20.2% mediated by AF
- The E_max_ of AF was reduced by 14.0% mediated by PQP
- The E_max_ of PQP was increased by 31.6% mediated by AF

Hence, an asymmetric interaction was quantified for the AF-PQP combination, as both, increases and decreases of the PD parameters EC_50_ and E_max_ were observed. The model predictions agreed well with the observed parasitaemia (Figure 2).

#### Artefenomel (AF) and ferroquine (FQ) combination

To corroborate the previous results a second antimalarial combination, with accessible clinical data in patients, was studied with the same approach. For the *in vitro* checkerboard-PRR interaction assay, concentrations of 6, 9, and 100 nM or 0, 6, 7, 8, 10, and 50 nM were chosen for AF and FQ, respectively. Simulations with the PD interaction parameters estimated from the *in vitro* checkerboard-PRR interaction assay were carried out as done before and compared with the clinical trial observations. The parameter estimates of the PK/PD model are presented in Table S4. Assuming Bliss Independence, the following PD interactions were estimated:

- The EC_50_ of FQ was reduced by 68.9% by AF.
- The E_max_ of AF was reduced by 46.9% by FQ.

Hence, the PD interaction was dependent on the concentration. At low concentrations around the EC_50_, a synergistic effect was quantified where the EC_50_ of FQ was reduced by AF. Yet, at higher concentrations, when E_max_ was reached, the E_max_ was limited to FQ and the higher E_max_ of AF was not attained. This can be seen in Figure 3 where the slope of AF is steeper at the highest concentration studied in mono as compared to the combination scenarios. The model predictions agreed well with the observed parasitaemia (Figure 3).

**Figure 3:**
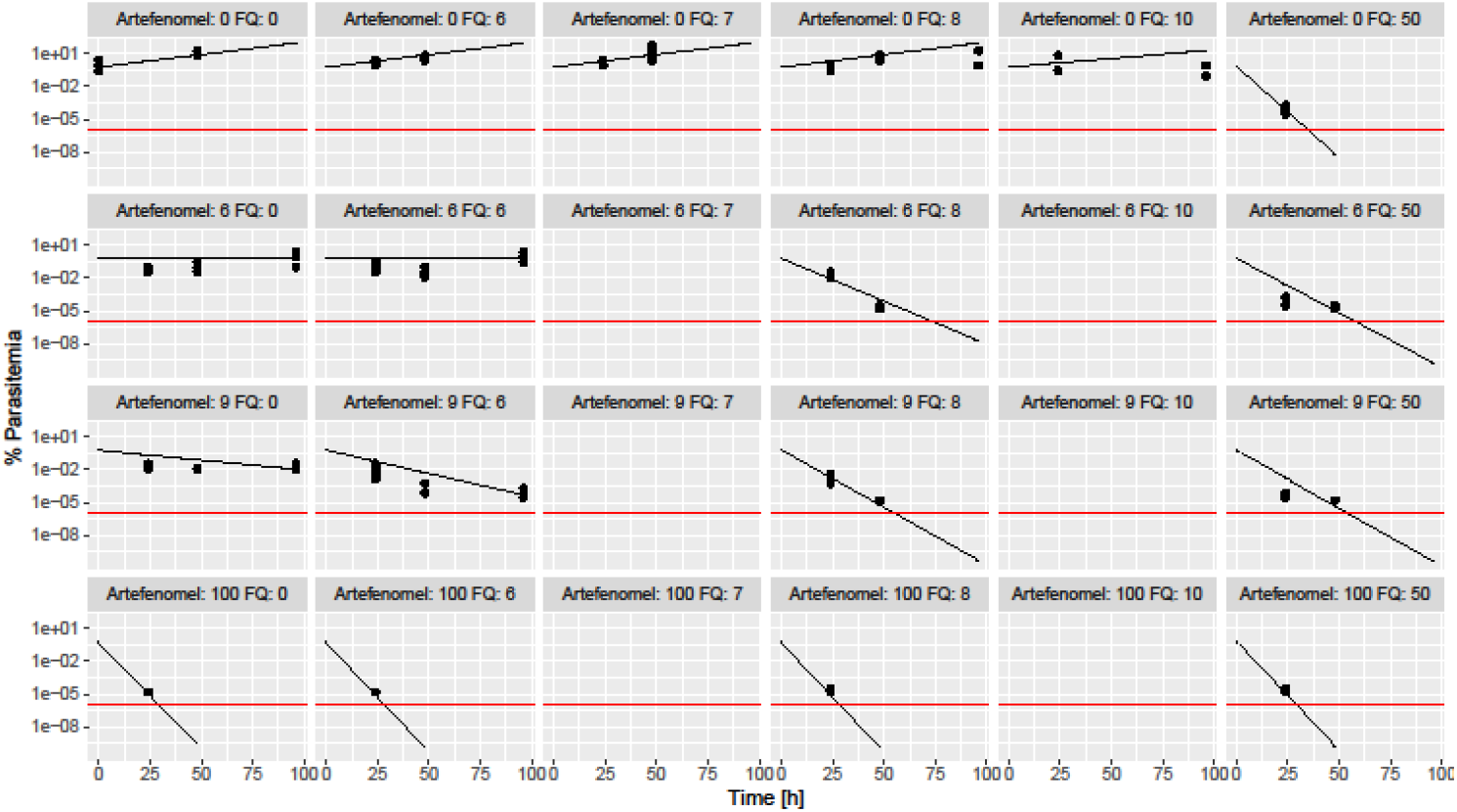
Individual model fits for artefenomel - ferroquine (AF - FQ) for the *in vitro* PRR checkerboard-PRR interaction assay data. Observed parasitaemia (black points) and model-predicted parasitaemia (black lines) time courses at different concentrations of AF and FQ (numbers in facet denote the concentration in nM of each combination partner); red line: quantification limit.

### *In vitro* checkerboard-PRR interaction assay estimates of PD interactions predict parasitological endpoints in humans equally well as *in vivo* animal models

To validate this new assay and the PD model, we used the PD interaction values (on EC_50_ and on E_max_) obtained *in vitro* to predict the parasitological endpoints of Phase 2 clinical trials. Furthermore, we compared the predictions obtained from our *in vitro* assay with the clinical data from patients (Figure 4A, B). For this purpose, a clinical pharmacometric model was developed that comprised a population pharmacokinetic submodel of the compounds and a pharmacodynamic submodel parameterized with the E_max_, EC_50_, Hill factor estimated with clinical monotherapy data and GPDI interaction estimates stemming from the new *in vitro* assay. The parasitological endpoints, adequate parasitological response at day 28 (APR28) and early treatment failure (ETF), calculated from the simulated profiles were compared with those calculated from the observed parasitaemia in patients. For both combinations, the *in vitro* checkerboard-PRR interaction model described the patients’ parasitological endpoints for most dose groups adequately well (Figure 4A, B). The scenarios of AF-PQP were all well predicted. For AF-FQ, underprediction was observed for APR28 at the lowest dose groups of AF. Yet, in none of the scenarios for both drugs, a significant difference between observed and predicted endpoint was observed (Figure 4).

**Figure 4:**
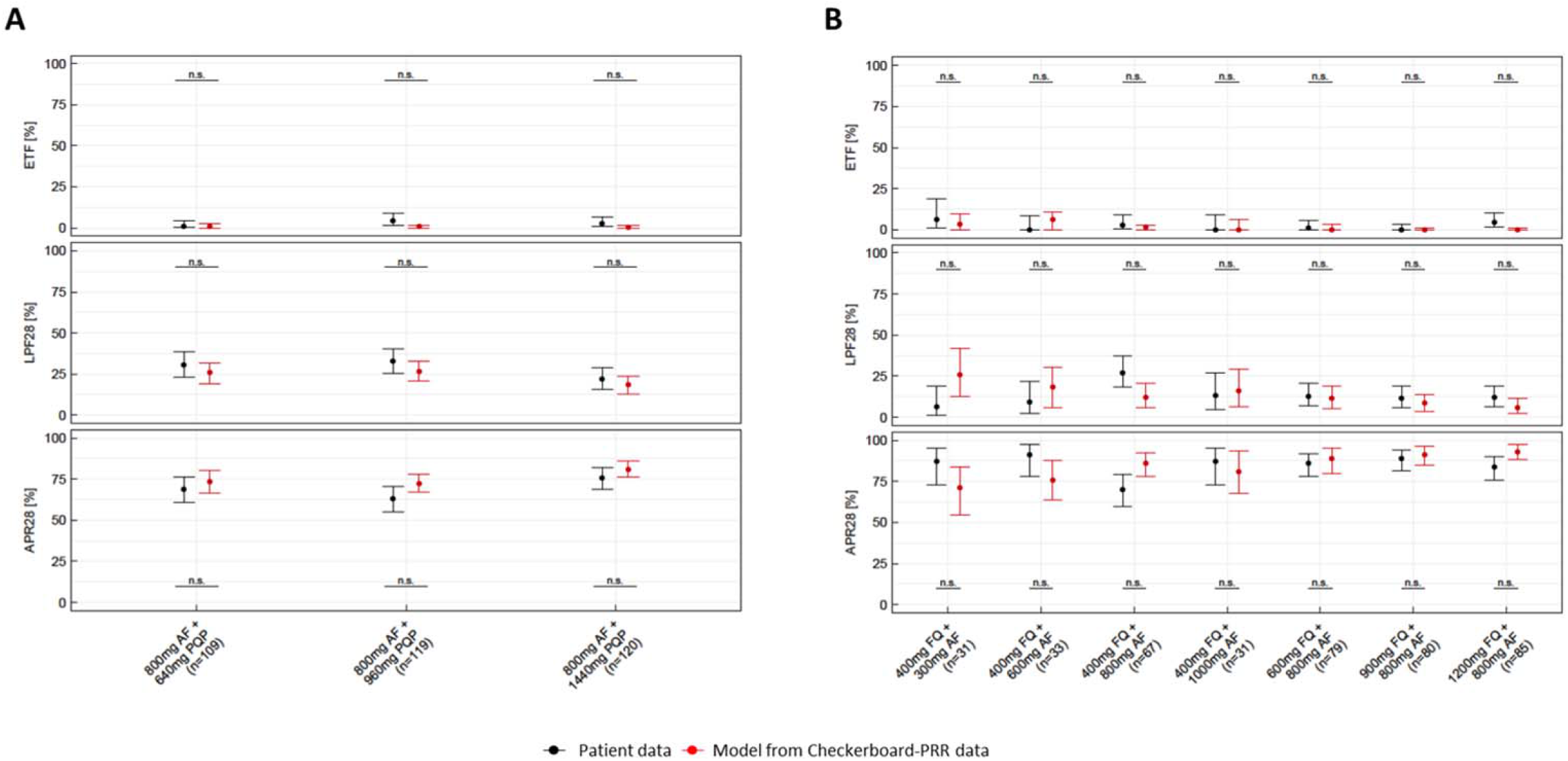
Comparison of parasitological endpoints calculated from observed and predicted parasitaemia in the patients receiving (A) AF - PQP or (B) AF - FQ. The parasitological endpoints: early treatment failure (ETF) and adequate parasitological response at day 28 (APR28) are summarized across for all patients per treatment group with of AF and PQP. Error bars depict 90% confidence intervals derived non-parametrically and by Clopper-Pearson for simulations and patients’ data respectively. Colour codes for patients’ data (black), and simulations with interaction parameters derived from *in vitro* checkerboard-PRR interaction assay (red).

In addition, data from previous studies for the AF - PQP obtained from *in vivo Pfalc*HuEryMouse and VIS combination studies, was compared to the *in vitro* checkerboard-PRR interaction assays (Figure S3, Table S5). We observed that PD interaction parameters derived from the *in vitro* checkerboard-PRR interaction assay or the *Pfalc*HuEryMouse model both adequately predicted the parasitological endpoints in patients. Of note, the GPDI model parameters from the *Pfalc*HuEryMouse model led to higher uncertainty in the model parameter estimates – likely due to sparser underlying data – and hence the prediction intervals were larger. The lowest agreement between observed parasitological endpoints in patients and corresponding model predictions was obtained when the VIS-derived PD interaction parameters were used. Parameter estimates from the *Pfalc*HuEryMouse and VIS are presented in Tables S2 and S3, respectively.

In conclusion we observed that the *in vitro* checkerboard-PRR interaction model described the patients’ parasitological endpoints for most dose groups adequately well. Furthermore, the *in vitro* checkerboard-PRR interaction assay allowed to estimate more complex interaction models including non-symmetric interactions with distinct perpetrator and victims within the PD interaction, as well as interactions on the EC_50_ as well as E_max_ level (Table S5). While the *in vivo* systems only allowed to estimate either EC_50_-based and/or E_max_-based interactions and the quantified interaction parameters were mutual. This underlines the value of the more granular PD data obtained from the *in vitro* PRR checkerboard as compared to the *in vivo* systems *Pfalc*HuEryMouse and VIS.

## DISCUSSION

In the present study experimental and-*in* silico-based approaches were combined to generate informative *in vitro* preclinical data to provide quantitative insights into PD interactions of antimalarial drugs and their translation into the clinical setting. This new *in vitro* combination assay leverages the previously developed gold standard PRR assay *(17)* to evaluate the maximum parasite killing rate by informing about parasite viability at different time-points after drug exposure and allows not only to characterize single drug effects but also to study drug combinations. Thereby the checkerboard-PRR interaction assay data was quantitatively evaluated using the GPDI model, which provided pharmacological insights into the observed drug interactions, *i*.*e*. shifts of EC_50_ and E_max_ in comparison to their single drug effects and the expected additive effect of the combination partners. Moreover, the *in vitro*-derived interaction parameters were used in clinical trial simulations. With the case examples of AF - PQP, we showed that the *in vitro-*derived parameters led to similar predictions as the interaction parameters derived from the *Pfalc*HuEryMouse *in vivo* model (Figure S3 and Table S2). Interestingly, the PD interaction estimates from VIS predicted the parasitological endpoints less accurately (Figure S3 and Table S3). This novel *in vitro* assay displays several key properties that advance historically used techniques in preclinical malaria research. Indeed, conventional *in vitro* assays used to study PD interaction, solely measure growth inhibition, and evaluate if the combined growth inhibition deviates from expected additive growth inhibition, and thus do not inform about PD killing interactions. While conventional assays thereby might provide some insight on potency changes in combination, no information on potential interactions of parasite killing (*i*.*e*. interactions on E_max_) can be obtained. Another widely used technique to study DDIs is the *Pfalc*HuEryMouse *in vivo* model. This model can provide quantitative insights into parasite killing, parasite resistance as well as PD drug interactions and thereby also provides valuable information that can be leveraged in a model-based approach for clinical prediction. Yet, this *in vivo* model is time-consuming, very costly and requires the use of chimeric mice which can pose ethically problems and – from a modelling perspective – provides less granular data compared to the *in vitro* assay.

The combined experimental-*in silico*-based approach displays several strengths: the development of an *in vitro* assay, in accordance with the 3R’s principle, that provides more granular raw data, and which is advantageous to explore the combined concentration-response surface in more detail than the previous models. The use of informative concentrations, based on effective concentrations *(16)* limits the number of scenarios to be tested to a reasonable number. The required time, workload as well as total cost of the *in vitro* PRR assay is far below the cost of animal experiments. Hence, this novel assay has the potential to streamline the drug development process to select and prioritize combination experiments. The data generated from the assay is used in conjunction with state-of-the art PK/PD modelling techniques. These models can describe the killing rate of the drugs over time alone and in combination. Thereby, the model not only provides interpretable estimates of the interaction (*i*.*e*. shifts of EC_50_ or E_max_), but it also allows to identify perpetrators or victims of the pharmacodynamic interactions *(18)*. In addition, since the model can describe the parasite killing rate over time, this *in vitro*-derived model can be linked to a clinical PK model of the drugs (using either first in human or predicted human PK data). This hybrid clinical PK-*in vitro* PD model can then be exploited for clinical trial simulations to evaluate the clinical potential of the combinations by predicting the doses in combination. This will allow the stratification and classification of different anti-malarial combinations treatments, allowing to select only those that are efficacious against drug-resistant strains and provide cure within a reasonable time (three days or less). The presented approach suggests that animal experiments can be reduced and performed in a confirmatory fashion, thus reducing the number of animals used. Yet, further research with additional matched case examples will be useful to further corroborate our findings. Few limitations and perspectives for further development shall be acknowledged. In contrast to *in vivo* experiments, the checkerboard-PRR interaction assay does not account for the PK profile of the tested drugs. While this was integrated in the simulations, studying constant concentrations does not provide any insight into persistent drug effects, *e*.*g*. an ongoing killing or inhibition of growth after removal of the drug. Moreover, drug degradation could not be included in the presented study. In future studies, a combination of the checkerboard-PRR interaction assay with hollow-fibre type experiments to mimic the PK of single and combination regimens will be evaluated *(19, 20)*. Lastly, although the PRR readout represents the current gold-standard to quantify parasite killing, the assay is time intense as at very low parasitaemia levels up to 21 days are needed to quantify the parasitaemia. Alternative techniques such as the MitoTracker *(21, 22, 23)* or immuno-enzymatic assays measuring proteins such as *Plasmodium falciparum* lactate dehydrogenase enzyme or the histidine rich protein-2 *(24, 25)* should be explored to see whether these can provide a comparable, but less time-consuming readout to measure parasite killing. In conclusion, the implementation of this novel alternative translational technology as a “routine” *in vitro* screening process early in the drug discovery process will facilitate the gathering of more accurate data and improve the quality of preclinical models used to inform first-in-human clinical studies about potential DDIs. With the combination of highly advanced modelling and simulation techniques, the *in vitro*-derived interaction parameters provided similar predictions as those obtained from the more complex *in vivo* mouse model.

Moreover, in the presented study we showed that this novel *in vitro* assay can provide detailed information of PD drug interactions of antimalarials allowing stratification and classification of new antimalarial combinations, and potentially in other therapeutic areas. Thus, it helps to optimally select only the most promising combinations and doses for the clinical setting early in the drug development process, and thereby significantly reduces the number of animals conventionally needed in the preclinical phase while also hoping to minimize the attrition rate in clinical trials.

## MATERIALS AND METHODS

### Monotherapy parasite growth inhibition assay (pre-test)

Concentrations to be tested in the checkerboard-PRR are defined from ‘inhibitory concentrations’ in a pre-test assay to avoid parasite overgrowth in the PRR step. Briefly, in a 96-well plate, six serial dilutions (1:2) are performed of the desired working dilutions. Infected red blood cells (iRBCs; parasitaemia 0.3%, haematocrit 2.5%) are added on top of the compounds (1:2 dilution, 200 μL final volume per well). Assay plates are incubated at 37°C at 93% N_2_, 4% CO_2_, and 3% O_2_. Parasite growth following drug treatment is quantified via the incorporation of [^3^H] hypoxanthine as previously described *(26)*. After 48 hours, [^3^H] hypoxanthine (= 0.25μCi) solution is added to each well and the plates are incubated for another 24 hours. Plates are harvested with a Betaplate™ cell harvester (Wallac, Zurich, Switzerland), which transfers the lysed red blood cells onto a glass fiber filter. The dried filters are inserted into a plastic foil with 10 mL of scintillation fluid and counted in a Betaplate™ liquid scintillation counter (Wallac, Zurich, Switzerland). The results are recorded as counts per minute (cpm) per well at each compound concentration. Data are transferred into a graphic programme (*e*.*g*. EXCEL, GraphPad Prism) and expressed as percentage of the untreated controls.

### Determination of single drug effects from the growth inhibition assay

In the first step, the *IC*_50_ is calculated from conventional growth inhibition data using the sigmoidal maximum effect model with nonlinear regression, relating the observed growth inhibition *I* to the drug concentration *C*:

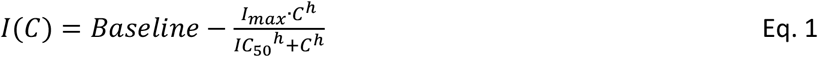

where *Baseline* represents the observed readout of the growth control, *I*_*max*_ the maximum reduction of the readout compared to *Baseline, IC*_50_ the concentration *C* displaying half-maximal reduction of the readout and the hill coefficient *h* the steepness of the concentration-effect relationship.

The concentrations for the checkerboard-PRR interaction assay were chosen from the pre-test so that at least three informative concentrations were selected covering the *IC*_20_ (*i*.*e*. the concentration stimulating 20% of the of the maximum effect), *IC*_80_ (*i*.*e*. the concentration simulating 80% of the maximum effect) and a ‘high’ concentration covering the clinically relevant maximum concentrations, adapted from Chen *et al. (16)*.

### Checkerboard-parasite reduction ratio (PRR) interaction assay

The newly developed *in vitro* checkerboard-PRR interaction assay is based on the original PRR assay *(17). P. falciparum* NF54 drug-sensitive parasite strain was cultivated in RPMI 1640 medium supplemented with 0.5% ALBUMAX® II, 25 mM Hepes, 25 mM NaHCO_3_ (pH 7.3), 0.36 mM hypoxanthine, and 100 μg/mL neomycin, as previously described *(27, 28)*. Cultures were maintained in an atmosphere of 3% O_2_, 4% CO_2_, and 93% N_2_ and at 5% haematocrit in humidified modular chambers at 37°C. To initiate the assay, the parasite culture was adjusted to 0.5% parasitaemia and 2% haematocrit, and 5 mL of the parasite culture were distributed per well into 6-well plates. Compound powders were dissolved in dimethyl sulfoxide (DMSO) to obtain 10 mM stocks. Thereafter, compounds and compound combinations were prepared in hypoxanthine-free medium and added to the corresponding well to achieve the desired concentrations. An untreated control was included to monitor parasite growth up to 48 hours. Cultures were incubated in an incubation chamber as described above. After 24, 48, 72 and/or 96 hours, 1 mL of culture was sampled from each well, the compound was removed by washing twice before resuspending the blood pellet and serially diluting it in 96-well plates. For each time point and each compound or compound combination, 15 serial dilutions of four technical replicates were performed and afterwards plates were incubated as described above (Figure S1). Medium was refreshed once a week and fresh erythrocytes were provided to allow growth of parasites. After 18-19 days, the culture medium was removed and replaced with screening medium containing 0.5 μCi of [^3^H] hypoxanthine as previously described *(27, 28)*. Another 48-72 hours later, plates were frozen at -20°C for a minimum of 24 hours. Thawed plates were harvested with a Betaplate™ cell harvester onto glass-fiber filters. The radioactivity was quantified using a Betaplate liquid scintillation counter and the results were expressed as counts per minute (cpm) per well. In addition, coloured spots on the filter mats were recorded. They served, together with observed medium colour changes during the growth period as visual indicators of parasite growth. Pyrimethamine at 10 times the *IC*_50_ value served as an internal control and was sampled at 24, 48, 72, and 96 hours to obtain a full-time-course viability curve. For each technical replicate of a sample, the number of viable parasites was extrapolated using the following equation:

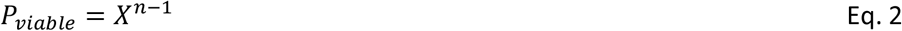

where *P*_*viable*_ represents the number of viable parasites, *X* the dilution factor used for serial dilution and *n* the number of wells rendering parasite growth. This number was then used to determine the well-specific parasitaemia.

The efficacy studies in *P. falciparum* infected NGS mice as well as the human volunteer infection study to develop the pharmacometric models to be compared with the estimates and predictions derived from the *in vitro* checkerboard PRR assay are presented in the Supplementary Material.

### Patients’ infection study

The human patient’s data used in this study were obtained from randomised, single-dose, Phase 2b clinical trials as previously described *(4)(3)*. Briefly, male and female African or Asian patients aged ≥6 months to <70 years with uncomplicated *P. falciparum* malaria and with microscopically confirmed (blood smear) *P. falciparum* mono-infection were treated with AF and FQ *(3)* or AF and PQP *(29)*. Patients were hospitalized for a minimum of 48 h or 72 h post-dose. Following discharge from the hospital, patients returned for further assessments at different days post treatment up to 63 days to assess treatment efficacy, safety, and PK. Parasitaemia was monitored by microscopy and/or PCR. The studies were conducted in accordance with consensus ethics principles derived from international ethics guidelines, including the Declaration of Helsinki, and the International Council for Harmonisation of Technical Requirements for Pharmaceuticals for Human Use guidelines for Good Clinical Practice, all applicable laws, rules, and regulations. They were approved by the relevant Independent Ethics Committees (IECs) and, where relevant, local regulatory authorities at each of the participating sites. The trials were registered in the ClinicalTrials.gov registry with registration no. NCT02497612 for AF and FQ and NCT02083380 for AF and PQP.

#### Modelling of PD

All *in vitro* checkerboard-PRR interaction assay data was analysed using non-linear regression analysis in the NONMEM® software (ICON development service, Gaithersburg, MD, version 7.4), while assuming constant concentrations.

In the first step, the parasite growth parameters were estimated with an exponential growth model using viable parasites from the checkerboard-PRR interaction assay at 48 and 96 hr. The growth model is defined as follows:

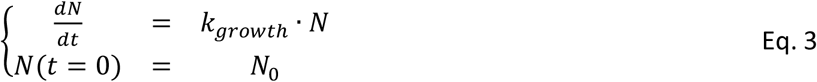

Where *N*(*t*) represents the model-predicted viable parasites at time *t* and *k*_*growth*_ the first-order growth rate and *N*_0_ the initial condition. *N*_0_ was estimated from the back-extrapolated raw readout at *t =* 0hr. The raw read *Y* was related to *N* by a proportional residual error model and a *Baseline*, which was estimated from the raw readout of the negative controls of the checkerboard, as follow:

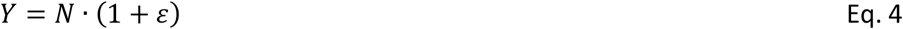

Subsequently, the mono drug effects were estimated from the wells containing only single drug effects using a sigmoidal maximum effect model:

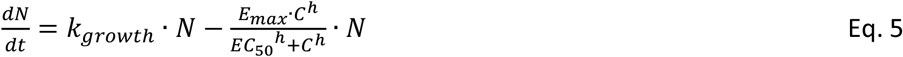

where *E*_*max*_ represents the maximum kill rate, *EC*_50_ the concentration stimulating half-maximum kill and the hill coefficient *h* the steepness of the concentration-effect relationship.

Thereafter, the combined drug effects for the drugs ‘1’ and ‘2’ are evaluated. First, the predicted additivity is computed assuming no drug interaction under bliss independence criteria fixing all model parameters.

Bliss Independence:

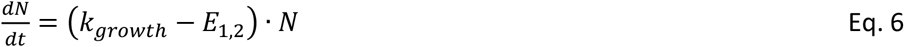

where:

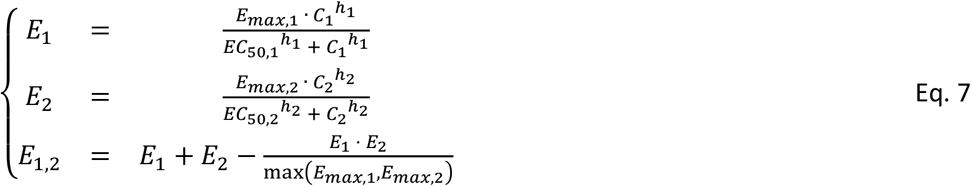

The model fit is evaluated by computing the Akaike Information Criterion (AIC). A lower AIC indicates a better model fit for the additivity model, which is then taken forward in the analysis. In case one of the drugs displayed a lag phase, a first order time delay rate constant 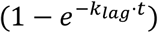 was added on the respective killing rate.

The PD interactions are evaluated using the General PD Interaction (GPDI) model *(18)* as shifts of *EC*_*max*_ and *EC*_50_ here exemplified for EC50as shown in Eqs. 8 and 9 as:

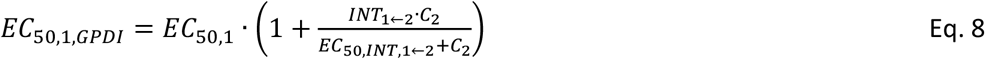

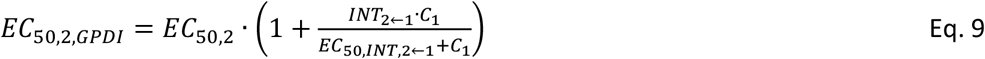

where *EC*_50,1,*GPDI*_ represents the *EC*_50_ of drug 1 being shifted by drug 2 at maximum by *INT*_1←2_ at a potency of *EC*_50,*INT*,1←2_, and *vice versa* for *EC*_50,2, *GPDI*_. The GPDI model is build-up using statistical criteria (likelihood ratio test, alpha of 0.05, *df = 1*, all model parameters estimated) starting with a reduced model with a single *INT* parameter for each drug indicating mono-directional interactions, up to estimating both *INT* parameters and finally also estimating the interaction potencies (*EC*_50,*INT,i*←*j*_).

An estimated *INT* parameter between -1 and 0 indicates that the respective *EC*_50_ is decreased in presence of the other drug, demonstrating synergy. An estimated *INT* parameter not significantly different from zero indicates additivity. An estimated *INT* parameter >0 indicates that the *EC*_50_ is increased in presence of the other drug, demonstrating antagonism. *INT* parameters of opposite polarity indicate asymmetric interactions where overall antagonism or synergy cannot be concluded, and interactions are concentration dependent.

### Clinical trial simulations and comparison using *Pfalc*HuEryMouse, VIS and *in vitro* checkerboard-PRR interaction assay estimates

The ability to predict the outcome of combination therapies in humans (VIS and/or clinical trials) using *Pfalc*HuEryMouse, VIS, and *in vitro* checkerboard-PRR interaction parameter estimates was assessed in clinical trial simulations of two anti-malarial combo treatments. Treatment effects in individual patients were simulated using interaction parameters from *in vitro* checkerboard-PRR interaction assay, *Pfalc*HuEryMouse-model experiments and VIS studies and compared to clinical study data. For each subject in the clinical trial, parasitaemia was simulated using individual PK parameters, baseline parasitaemia (*PL*_*base*_) and parasite growth rate; respective monotherapy PKPD models and drug PD interaction estimates from *Pfalc*HuEryMouse, VIS or *in vitro* checkerboard-PRR. Uncertainty in the PD parameters were incorporated by simulating 250 trials. For each trial, interaction parameters were sampled from uncertainty, described by the relative standard error, based on estimation results from *in vitro* checkerboard-PRR, *Pfalc*HuEryMouse model and VIS. Furthermore, for each trial, *E*_*max*_, *EC*_50_, *h* and the standard deviation (SD) for inter-individual variability (IIV) were sampled from uncertainty. Individual *E*_*max*_, *EC*_50_ and *h* were then drawn from IIV for each individual within a trial. The parasitaemia profile was calculated from the PK/PD model for each individual. A cure threshold of 1 parasite per body which corresponds to 1 parasite per 70 mL of blood per kg of bodyweight was defined. Once parasitaemia reached this threshold the parasitaemia level was set to the cure threshold for the remaining simulation time to indicate a complete eradication of parasites. Then, for each subject the following endpoints were calculated.

1. Early treatment failure (ETF): parasitaemia on day 2 post treatment was higher than on day 0 or if parasitaemia on day 3 was larger than 25 % of day 0 level.
2. Late parasitological failure at day 28 (LPF28): parasitaemia above the lower limit of quantification (LLOQ) on any day between day 7 and 28 in patients who did not meet criteria of the ETF.
3. Absence of parasitaemia at day 28 (APR28): no parasitaemia above LLOQ on day 28 in patients who did not meet any criteria of ETF or LPF28.

Simulated parasitological endpoints for each dose group were reported as median with 90% CI of the fraction of individuals achieving an endpoint per trial over all trials. A graphical outline of the simulation workflow is presented in Figure S2. Simulations and patient data were compared using Kolmogorov-Smirnov test. Due to the binary output for the endpoints, the fraction of individuals achieving an endpoint per trial were discrete. For statistical tests, the output of the simulations was smoothened using the bandwidth of the kernel density estimate. Resulting negative values of this smoothening procedure were set to 0. P-values were adjusted for multiple testing using the Benjamini-Hochberg procedure for each model *(30)*.

Of note, the parasitological endpoints were calculated for VIS subjects from the observations and predictions only to assess how the PD interaction parameters performed. To extrapolate clinical outcomes from a VIS, simulations are run with baseline parasitaemia sampled from a clinical dataset rather than VIS subjects: baseline parasitaemia are higher in patients and for a same dose level, fewer patients will succeed the clinical criterion than VIS subjects.

## Acknowledgments

We acknowledge our colleagues at Medicines for Malaria Venture, our collaborators at the Swiss TPH Parasite Chemotherapy Unit, Iñigo Angulo Barturen and María Belén Jiménez-Díaz from The Art of Discovery (TAD) for the *in vivo* animal biological data.

## Author contributions

Conceptualization: CDG, SW and MR

Methodology: CDG, SW and MR

Investigation: CDG, SW and MR

Visualization: SW

Study design: CDG, SW, MR and AW

Experimental work: AW, CG

Data analysis: SW, MHCR, NB, KK and NG

Data interpretation: CDG, SW, MHCR, NB, NG and MR

Writing-original manuscript: CDG and SW

Writing-review and editing: CDG, SW, MR, MHCR, NB, AW, JM and NG

Oversight, acquisition of funding and key experimental materials: CDG, SW and MR

## Data and materials availability

The datasets analysed during the current study are available from the corresponding author on request and with permission of Medicines for Malaria Venture.

